# First nuclear genome assembly of an extinct moa species, the little bush moa (*Anomalopteryx didiformis*)

**DOI:** 10.1101/262816

**Authors:** Alison Cloutier, Timothy B. Sackton, Phil Grayson, Scott V. Edwards, Allan J. Baker

## Abstract

High throughput sequencing (HTS) has revolutionized the field of ancient DNA (aDNA) by facilitating recovery of nuclear DNA for greater inference of evolutionary processes in extinct species than is possible from mitochondrial DNA alone. We used HTS to obtain ancient DNA from the little bush moa (*Anomalopteryx didiformis*), one of the iconic species of large, flightless birds that became extinct following human settlement of New Zealand in the 13 ^th^ century. In addition to a complete mitochondrial genome at 249.9X depth of coverage, we recover almost 900 Mb of the moa nuclear genome by mapping reads to a high quality reference genome for the emu (*Dromaius novaehollandiae*). This first nuclear genome assembly for moa covers approximately 75% of the 1.2 Gb emu reference with sequence contiguity sufficient to identify more than 85% of bird universal single-copy orthologs. From this assembly, we isolate 40 polymorphic microsatellites to serve as a community resource for future population-level studies in moa. We also compile data for a suite of candidate genes associated with vertebrate limb development and show that the wingless moa phenotype is likely not attributable to gene loss or pseudogenization among this candidate set. We also identify potential function-altering coding sequence variants in moa for future experimental assays.

## Introduction

The extinct moa of New Zealand (Aves: Dinornithiformes) comprise nine currently recognized species (Bunce et al. 2009) and belong to the Paleognathae, which encompasses the flightless ratites (ostrich, emu, cassowary, kiwi, rheas, moa, and elephant birds) and the volant, or flying, tinamous. Extinction of all moa species is thought to have closely followed Polynesian settlement of New Zealand in the late 13 ^th^ century as the result of direct human exploitation compounded by anthropogenic land-use changes and impacts associated with introduced species (Allentoft et al. 2014; Holdaway et al. 2014).

In addition to a rich history of paleontological study (reviewed in Worthy and Holdaway 2002), ancient DNA has yielded multiple novel insights into moa biology (Allentoft and Rawlence 2012; Grealy et al. 2017). Cooper et al. (1992) first used aDNA amplified by the polymerase chain reaction (PCR) to show that moa are not most closely related to kiwi, indicating independent arrivals of these two lineages to New Zealand. Instead, aDNA places moa as sister to the volant tinamous, consistent with multiple independent losses of flight in ratites (Phillips et al. 2010; Haddrath and Baker 2012; Baker et al. 2014; Cloutier et al. 2019; Sackton et al. 2019). Ancient DNA has also helped clarify moa taxonomy (Baker et al. 2005; Bunce et al. 2009), and was instrumental in identifying extreme reversed sexual size dimorphism that misled some morphological taxonomic designations (Bunce et al. 2003; Huynen et al. 2003). Contributions from ancient DNA have ‘clothed’ moa by assigning feathers to their species of origin (Rawlence et al. 2009), identified males as the likely incubating sex from eggshell aDNA (Huynen et al. 2010), and investigated moa feeding ecology and parasites using coprolites (Wood et al. 2013a,b).

This diversity of aDNA research testifies to the wealth of relatively well-preserved moa remains (Allentoft and Rawlence 2012). Yet, most molecular studies of moa have relied upon mitochondrial DNA (mtDNA) since mtDNA occurs in high copy number per cell and is therefore more readily recovered than nuclear DNA from subfossil substrates, where aDNA is often highly degraded (Allentoft and Rawlence 2012; Hofreiter et al. 2015; Grealy et al. 2017). High throughput sequencing (HTS) has revolutionized the field of aDNA by allowing recovery of these short segments of nuclear DNA. Unlike mtDNA, which is uniparentally inherited and represents only a tiny fraction of the total genomic ‘blueprint’ in an individual, nuclear DNA can provide much more detail about the evolutionary history and unique adaptations of extinct species (Hofreiter et al. 2015; Grealy et al. 2017). It is therefore likely that we have only just begun to access the available genetic information for moa.

We used high throughput sequencing to recover aDNA from the little bush moa *(Anomalopteryx didiformis).* Little bush moa were distributed in lowland forests across the North and South Islands of New Zealand and were among the smallest of moa species, reaching heights of 50–90 centimeters (Worthy and Holdaway 2002; Bunce et al. 2009). In addition to a complete mitochondrial genome, we report the first nuclear genome for any moa species, assembled by mapping little bush moa reads to a high quality draft genome for the emu *(Dromaius novaehollandiae,* Sackton et al. 2019). We use this moa nuclear genome to isolate polymorphic microsatellites for future population-level studies and also recover coding sequence for a suite of candidate genes to investigate their possible association with flightlessness in moa and other ratites.

## Materials and Methods

### DNA extraction and sequencing

DNA was extracted from a single toe bone of an unprovenanced moa specimen held in the collections of the Royal Ontario Museum (ROM; Toronto, Canada). Some of the HTS reads reported here were previously used for phylogenetic analysis of paleognath relationships using 1,448 ultraconserved elements (UCEs) and protein-coding loci (Baker et al. 2014), and PCR-based sequences obtained from this same specimen have been reported by Haddrath and Baker (2001), Baker et al. (2005, sample A. did. OH), and Haddrath and Baker (2012, sample TW95).

DNA extraction followed Baker et al. (2005). In brief, the outer 1–2 mm was removed from the bone surface by microblasting with an Airbrasive System (MicroBlaster; Comco, Burbank CA, USA), and 0.2 grams of the remaining material was ground into fine powder. Enzymatic digestion proceeded overnight at 56°C in buffer containing final concentrations of 0.5M EDTA, 200 μg/mL proteinase K, and 0.5% N-laurylsarcosine at pH 8.0 (Hagelberg 1994), and DNA was purified using commercially available silica spin columns (DNeasy Blood & Tissue Kit; Qiagen, Germantown MD, USA). Sample preparation occurred in a dedicated aDNA workspace in the ROM following established best practices (Cooper and Poinar 2000; Knapp et al. 2011).

Library preparation and sequencing was performed by The Centre for Applied Genomics, The Hospital for Sick Children, Toronto, Canada. Library A_didi_CTTGTA was constructed from 200–400 bp size-selected DNA sheared to 200 bp insert size followed by library preparation with the Illumina TruSeq DNA v3 DNA Prep Kit. Paired-end sequencing (2 x 101 bp) was carried out on three lanes of a HiSeq 2500 platform using Illumina v3 chemistry. A second TruSeq library (A_didi_GCCAAT) was prepared from the same input DNA and sequenced on two partial lanes of a HiSeq 2500. Three additional libraries were constructed with the Illumina Nextera XT Sample Preparation Kit. The A_didi_CAGAGA and A_didi_CTCTCT libraries used input DNA < 500 bp with no additional shearing, while the A_didi_AGGCAG library used DNA 500 bp–2 kb in size subsequently sheared to < 700 bp. These Nextera libraries were pooled for sequencing on a single HiSeq 2500 lane.

### Read processing and genome assembly

Trimmomatic v. 0.32 (Bolger et al. 2014) was run in paired-end mode for adapter removal and quality trimming and reads with post-trimming length below 25 bp were discarded (options ILLUMINACLIP:[adapter_file]:2:30:10:1:true SLIDINGWINDOW:10:13 MINLEN:25).

A *de novo* mitochondrial genome assembly was built with MITObim v. 1.8 (Hahn et al. 2013) using the published little bush moa mtDNA genome as a starting seed (GenBank accession NC_002779, Haddrath and Baker 2001).

To obtain a nuclear sequence assembly, we first mapped reads to a draft genome for emu *(Dromaius novaehollandiae*; Sackton et al. 2019, GenBank accession GCA_003342905.1), and then re-mapped reads to the initial moa consensus for improved recovery of short and/or variant reads. First, to estimate an appropriate substitution parameter between the emu and moa, a random subset of reads was mapped to the emu reference with Stampy v. 1.0.28 (Lunter and Goodson 2011) using default settings. The full data were then mapped to emu with Stampy and this user-specified substitution parameter (estimated at 0.0839). Samtools v. 1.3.1 (Li et al. 2009) was used to filter reads with mapping quality score below 30, and duplicates within each library were marked and removed with Picard Tools v. 2.6.0 (https://broadinstitute.github.io/picard/) before merging mapped reads across libraries. Samtools ‘mpileup’ was used to output variant call format (VCF) files with minimum mapping quality 30 and base quality 20, and a consensus sequence was called with BCFTools v. 1.2. Reads were remapped to this initial consensus with Bowtie2 v. 2.2.9 (Langmead and Salzberg 2012), with subsequent post-processing as above.

In addition to the nuclear assembly detailed above (hereafter referred to as the ‘original assembly’), an error-corrected version of the genome was also generated to account for nucleotide misincorporations due to post-mortem DNA damage characteristic of ancient DNA. Trimmed reads were processed with PEAR v. 0.9.7 (Zhang et al. 2014b) to merge overlapping read pairs before mapping to the original moa assembly using Bowtie, followed by filtering and duplicate removal as detailed above. The resulting BAM file was used as input to mapDamage2 v. 2.0.7 (Jónsson et al. 2013) with default parameters and the --rescale option specified to recalibrate base quality scores. Samtools v. 0.1.11 was used to generate ‘pileup’ files with minimum base quality 20 after mapDamage recalibration, and a custom Perl script (available in the accompanying Dryad Digital Repository [DOI pending]) was used to mask bases with no coverage following recalibration.

For both the original and mapDamage corrected assemblies, genome completeness was measured with BUSCO v. 2.0 and the aves_odb9 data set (Simão et al. 2015) to search for 4,915 bird universal single-copy orthologs. Mosdepth v. 0.2.5 (Pedersen and Quinlan 2018) was used to calculate per-site depth of coverage (DoC) while avoiding ‘double counting’ overlapping read pairs.

### Taxonomic read profiling

Trimmed reads were queried against a custom database containing all avian, bacterial, archaeal, plant (including algae and fungi), and viral sequences from GenBank Release 217 as well as publicly available genomes for the chicken *(Gallus gallus,* galGal4 release, Hillier et al. 2004), North Island brown kiwi *(Apteryx mantelli,* Le Duc et al. 2015), ostrich *(Struthio camelus,* Zhang et al. 2014a), white-throated tinamou *(Tinamus guttatus,* Zhang et al. 2014a), human (reference genome GRCh38), and the draft emu assembly (Sackton et al. 2019). Reads were mapped in BlastN mode with default parameters in MALT v. 0.3.8 (accessed from http://ab.inf.uni-tuebingen.de/data/software/malt/download/welcome.html), and MEGAN Community Edition v. 6.6.4 (Huson et al. 2016) was used for taxonomic clustering.

### Identification of polymorphic microsatellite repeats

MSATCOMMANDER v. 1.0.8 (Faircloth 2008) was used to identify all dinucleotide microsatellites with a minimum of six repeat units and all trinucleotides with at least four repeats in the moa nuclear assembly. Candidate loci with more than 10% uncalled bases (Ns) in the region encompassing the microsatellite and 250 bp of flanking sequence to either side were excluded. Reads mapped to each remaining candidate region were realigned using STR-realigner v. 0.1.01 (Kojima et al. 2016) and genotypes were called with RepeatSeq v. 0.8.2 (Highnam et al. 2013). Heterozygous loci with minimum genotype likelihood ≥ 10 and minimum depth of coverage ≥ 2 for both reference and alternate alleles were retained. Sequence for each retained locus (repeat + flank) was used in blastn searches against draft genomes for seven ratites from Sackton et al. (2019), and the ostrich (Zhang et al. 2014a). Blastn hits with evalue < 1e^-10^ were extracted from reference genomes and aligned with MAFFT v. 7.245 (Katoh and Standley 2013).

### Tests of selection for candidate limb development genes

Multiple sequence alignments were compiled for 26 candidate genes with established roles in vertebrate limb development (reviewed in Zakany and Duboule 2007; Tanaka 2013; Tickle 2015; Petit et al. 2017; listed in Table 2a) and for 11 genes with potential function-altering variants in the Galapagos cormorant *(Phalacrocorax harrisi*) hypothesized to accompany phenotypic modifications typical of flightless birds (Burga et al. 2017, listed in Table 2b). Gene models were manually curated for ten new draft genome assemblies for paleognaths (Sackton et al. 2019). Moa coding sequence was obtained from pairwise whole-scaffold alignments of moa to emu using reference emu coordinates (alignments and an accessory Perl script are available in Dryad). Sequences from draft paleognath genomes were combined with available avian sequences from GenBank and cormorant sequences from Burga et al. (2017). Amino acid translations were aligned with MAFFT v. 7.245 (Katoh and Standley 2013). Partial (< 70% of total alignment length) and poorly aligning (< 60% mean pairwise amino acid identity) sequences were removed, and the resulting alignment was used to guide gap insertion in the corresponding nucleotide sequences. GenBank source information, curated gene models, and sequence alignments are available in Dryad.

**Table 1.** Assembly statistics for emu reference and little bush moa nuclear genomes

|  | **Emu** | **Little bush moa** |  |
| --- | --- | --- | --- |
|  |  | **Original assembly** | **MapDamage corrected** |
| No. Scaffolds | 2,777 | 1,909 | 1,843 |
| Total scaffold length (bp, gapped) | 1,192,237,364 | 1,190,759,065 | 1,189,933,016 |
| Total ACGT bases (bp) | 1,179,039,690 | 889,719,452 | 868,145,842 |
| No. Contigs<sup>1</sup> | 18,796 | 1,335,633 | 1,368,477 |
| Contig N50 (bp) | 187,464 | 1,127 | 1,068 |
| Mean contig size (bp) | 62,805 | 673 | 643 |
| Avg. break between contigs (bp) | 733 | 218 | 226 |
| Total BUSCOs | 4797/4915 (97.6%) | 4299/4915 (87.5%) | 4190/4915 (85.2%) |
| Complete BUSCOs | 4629/4915 (94.2%) | 3701/4915 (75.3%) | 3550/4915 (72.2%) |

<sup>1</sup>Contig breaks are defined by strings of $\geq 25$ consecutive Ns

**Table 2.** Tests of selection for candidate limb development genes using moa sequence from the original genome assembly

| Gene | Description | CDS length<br>(AA, % of total) |  |  | RELAX tests |  |  |  |
| --- | --- | --- | --- | --- | --- | --- | --- | --- |
|  |  | Chicken | Emu | Moa | Moa |  | All flightless |  |
|  |  |  |  |  | K | P<sub>adj</sub> | K | P<sub>adj</sub> |
| a) Candidate limb development genes |  |  |  |  |  |  |  |  |
| FGF8 | Fibroblast growth factor 8 | 214 | 214 | 189 (88%) | 3.912 | 0.991 | 49.994 | 0.081 |
| FGF10 | Fibroblast growth factor 10 | 212 | 212 | 204 (96%) | 0.494 | 0.419 | 1.052 | 0.542 |
| GLI3 | GLI family zinc finger 3 | 1576 | 1575 | 1570 (99%) | 1.491 | 0.389 | 0.359 | 0.006 |
| HOXA1 | Homeobox A1 | 320 | 319 | 308 (96%) | 0.999 | 0.991 | 0.931 | 0.459 |
| HOXA2 | Homeobox A2 | 375 | 374 | 357 (95%) | 0.178 | 0.074 | 3.424 | < 0.001 |
| HOXA3 | Homeobox A3 | 413 | 413 | 414 (100%) | 1.491 | 0.389 | 1.033 | 0.525 |
| HOXA4 | Homeobox A4 | 309 | 145<sup>+</sup> | 155 (50%) | 0.365 | 0.389 | 2.564 | 0.006 |
| HOXA5 | Homeobox A5 | 270 | 270 | 251 (93%) | 29.348 | 0.289 | 0.633 | 0.215 |
| HOXA6 | Homeobox A6 | 231 | 231 | 231 (100%) | 0.828 | 0.808 | 0.272 | 0.365 |
| HOXA7 | Homeobox A7 | 219 | 219 | 219 (100%) | 28.556 | 0.389 | 1.156 | 0.425 |
| HOXA9 | Homeobox A9 | 260 | 261 | 249 (95%) | 1.985 | 0.389 | 2.040 | 0.084 |
| HOXA10 | Homeobox A10 | 364 | 317<sup>+</sup> | 289 (79%) | 1.239 | 0.934 | 2.914 | 0.062 |
| HOXA11 | Homeobox A11 | 297 | 297 | 255 (86%) | 0.303 | 0.389 | 1.453 | 0.110 |
| HOXA13 | Homeobox A13 | 290 | 290 | 269 (93%) | 1.179 | 0.934 | 0.858 | 0.459 |
| HOXD3 | Homeobox D3 | 413 | 248<sup>+</sup> | 247 (60%) | 1.211 | 0.808 | 1.326 | 0.425 |
| HOXD4 | Homeobox D4 | 237 | 237 | 202 (85%) | 1.125 | 0.966 | 9.667 | 0.006 |
| HOXD8 | Homeobox D8 | 268 | 147<sup>+</sup> | 146 (54%) | 0.607 | 0.497 | < 0.001 | 0.379 |
| HOXD9 | Homeobox D9 | 302 | 299 | 283 (94%) | 0.295 | 0.339 | 0.816 | 0.217 |
| HOXD10 | Homeobox D10 | 339 | 339 | 339 (100%) | 0.923 | 0.991 | 49.998 | 0.152 |
| HOXD11 | Homeobox D11 | 280 | 282 | 272 (96%) | 0.967 | 0.976 | 0.642 | 0.081 |
| HOXD12 | Homeobox D12 | 266 | 266 | 266 (100%) | 1.452 | 0.397 | 1.006 | 0.569 |
| HOXD13 | Homeobox D13 | 301 | 82<sup>+</sup> | 74 (25%) | 0.926 | 0.991 | 0.939 | 0.525 |
| SALL4 | Spalt-like transcription factor 4 | 1108 | 1111 | 1023 (92%) | 0.793 | 0.389 | 0.860 | 0.146 |
| SHH | Sonic hedgehog | 425 | 422 | 398 (94%) | 1.979 | 0.389 | 0.974 | 0.542 |
| TBX5 | T-box 5 | 521 | 538 | 426 (79%) | 0.240 | 0.389 | 1.021 | 0.545 |
| WNT2B | Wnt family member 2B | 385 | 330<sup>+</sup> | 260 (68%) | 0.636 | 0.339 | 1.703 | 0.081 |
| b) Candidate genes from the Galapagos cormorant |  |  |  |  |  |  |  |  |
| DCHS1 | Dachsous cadherin-related 1 | 3266 | 3267 | 3086 (94%) | 1.094 | 0.389 | 1.006 | 0.554 |
| DVL1 | Dishevelled segment polarity protein 1 | 712 | 655<sup>+</sup> | 633 (89%) | 1.326 | 0.397 | 0.814 | 0.127 |
| DYNC2H1 | Dynein cytoplasmic 2 heavy chain 1 | 4301 | 4295 | 3991 (93%) | 1.152 | 0.389 | 0.763 | 0.089 |
| EVC | EvC ciliary complex subunit 1 | 984 | 927<sup>+</sup> | 871 (89%) | 0.169 | 0.389 | 0.654 | 0.023 |
| FAT1 | FAT atypical cadherin 1 | 4645 | 4644 | 4489 (97%) | 1.493 | 0.013 | 0.880 | 0.024 |
| GLI2 | GLI family zinc finger 2 | 1528 | 1528 | 1528 (100%) | 0.408 | 0.013 | 0.927 | 0.262 |
| IFT122 | Intraflagellar transport 122 | 1245 | 1239 | 1202 (97%) | 0.035 | 0.808 | 1.311 | 0.146 |
| KIF7 | Kinesin family member 7 | 1412 | 1279<sup>+</sup> | 1226 (87%) | 0.683 | 0.389 | 1.204 | 0.146 |
| OFD1 | OFD1, centriole and centriolar satellite protein | 1012 | 1014 | 1002 (99%) | 0.978 | 0.991 | 0.006 | 0.102 |
| TALPID3 | KIAA0586 | 1523 | 1527 | 1474 (97%) | 0.748 | 0.389 | 0.035 | 0.023 |
| WDR34 | WD repeat domain 34 | 500 | 502 | 459 (91%) | 18.909 | 0.251 | 0.776 | 0.102 |

K: Relaxation parameter (values < 1 indicate relaxed selection on foreground branches, values > 1 denote intensified selection)
P<sub>adj</sub>: Adjusted P-value (Q-value) controlling for the false discovery rate at a significance level of 0.05 based on N= 37 genes tested
<sup>†</sup>Partial CDS recovered in the emu reference sequence

We used the adaptive branch-site random effects likelihood model (aBSREL, Smith et al. 2015) in HyPhy v. 2.3.3 (Kosakovsky Pond et al. 2005) to test for lineage-specific selection in moa. We used the RELAX method (Wertheim et al. 2015), also implemented in HyPhy, to test for changes in selection intensity along specified foreground branches. We first tested for changes in moa relative to other ratites by pruning the data set to contain only ratites and then setting moa as the foreground branch and other flightless ratite lineages as the background. Second, we assessed shifts in selection accompanying loss of flight by setting all flightless lineages as the foreground (including the Galapagos cormorant, penguins, ratites, and inferred flightless ancestors) and all volant lineages as the background. Multiple test correction to control the false discovery rate at 0.05 within each analysis used the qvalue package v. 2.2.0 (Storey 2015) to control for the number of candidate genes analyzed (N= 37) as well as a more conservative approach controlling for the expected genome-wide false discovery rate based on an estimated N= 16,255 homologous orthologous groups of genes (HOGs) in birds identified by Sackton et al. (2019).

Functional effects of moa sequence variants were assessed with PROVEAN v. 1.1.5 (Choi et al. 2012), using a threshold score < −5 to identify possible function-altering variants following Burga et al. (2017). We computed PROVEAN scores for moa substitutions relative to the emu reference sequence and additionally comparing moa to an inferred ancestral sequence for the common ancestor of moa and tinamous reconstructed in PAML v. 4.8 under the codon-based model (Yang 2007).

## Results and Discussion

### Library characterization and endogenous DNA content

HTS yielded 143.4 Gb of raw data (Supplementary Table S1, SRA accession SRP132423). Most data incorporated into the mitochondrial and nuclear genomes described below originated from library A_didi_CTTGTA (Fig. 1A), due in part to greater sequencing effort for this library. Library A_didi_GCCAAT produced fewer reads than expected and had a high level of sequence duplication due to suboptimal cluster density (Supplementary Table S1). Recovery of moa DNA from the three Nextera libraries was also limited, a result that could reflect the smaller amount of input DNA used in the Nextera protocol and/or a decreased amount of endogenous DNA in the size fractions assayed for these preparations.

**Fig. 1.**
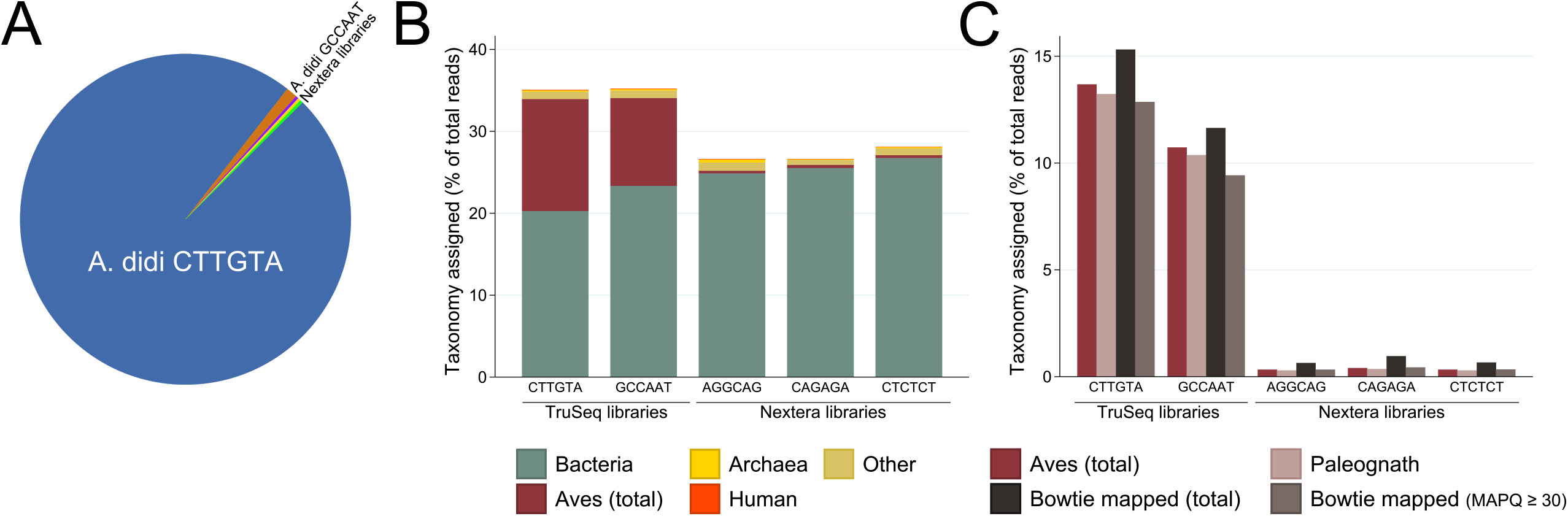
Endogenous DNA content across library preparations. A) Proportion of final nuclear genome assembly attributable to sequencing reads from each library. B) MEGAN assignment of taxonomic affiliations for individual sequencing reads. C) Proportions of reads assigned to avian species by taxonomic profiling compared to proportions mapped to the emu reference genome.

Taxonomic profiling of reads, which represent a mixture of endogenous moa aDNA and environmental DNA, assigned taxonomy to 26–35% of reads across libraries (Fig. 1B). The TruSeq libraries contained much higher proportions of reads assigned to Aves (e.g. all birds, 13% and 10% respectively for libraries CTTGTA and GCCAAT, Fig. 1B), with most of these reads further assigned to Paleognathae (Fig. 1C). Total mapping rates before duplicate removal mirror estimated amounts of endogenous DNA in each library (Supplementary Table S1, Fig. 1C), suggesting that use of a reference emu genome that was relatively divergent from moa for mapping nevertheless recovered most of the recognizably moa DNA within library extracts. Levels of read duplication (Supplementary Table S1) further indicate that sequencing saturation was reached to recover the maximum possible amount of endogenous DNA.

Ancient DNA is typically degraded to fragments smaller than 500 bp and displays characteristic post-mortem modifications leading to an excess of purines immediately preceding strand breaks and increasing cytosine (C) to thymine (T) substitutions toward fragment ends (Sawyer et al. 2012; Dabney et al. 2013). We cannot fully assess the extent of DNA damage because library construction involved DNA shearing, meaning that fragment ends represent a mixture of naturally occurring DNA breakage as well as strand breaks induced during library preparation. However, mean lengths of mapped reads and estimated insert sizes, especially for the two TruSeq libraries, are consistent with well-preserved DNA (Supplementary Table S1). Consequently, although we do observe signatures of aDNA damage, the amount of damage appears minimal (Supplementary Fig. S1). These observations are not unprecedented for well-preserved moa specimens. Cooper et al. (2001), Haddrath and Baker (2001), and Baker et al. (2005) successfully amplified moa PCR products 250–600 bp in length, and Cooper et al. (2001) reported high endogenous DNA content and little DNA damage for samples used to sequence complete mitochondrial genomes. Additionally, both the mitochondrial genome described below and phylogenetic analysis of genome-wide data sets of nuclear markers for this specimen corroborate its aDNA sequence authenticity (Baker et al. 2014; Cloutier et al. 2019; Sackton et al. 2019).

### Assembly of mitochondrial and nuclear genomes

We recovered a complete 17,043 bp mitochondrial genome at 249.9X average depth of coverage (DoC) following duplicate removal (Fig. 2A, GenBank accession pending). mapDamage correction yielded an identical mtDNA genome sequence. This new little bush moa assembly spans the entire 1,478 bp control region (D loop), which was not fully represented in the published mtDNA genome assembled from PCR-based sequencing of the same specimen (Haddrath and Baker 2001). The new HTS assembly is near identical to the existing reference, with only two SNPs across 777 bp of alignable control region sequence, and five SNPs and three single base pair indels across 15,565 bp lying outside the control region (99.9% identity), with all differences supported by > 50X DoC in the new HTS assembly. A 30 bp hypervariable control region ‘snippet’ diagnostic for moa lineages (McCallum et al. 2013) confirms taxonomic assignment of the sequenced specimen, and a longer (382 bp) segment spanning this region is identical to a haplotype from little bush moa sampled at multiple sites across the South Island of New Zealand (Supplementary Fig. S2, Bunce et al. 2009).

**Fig. 2.**
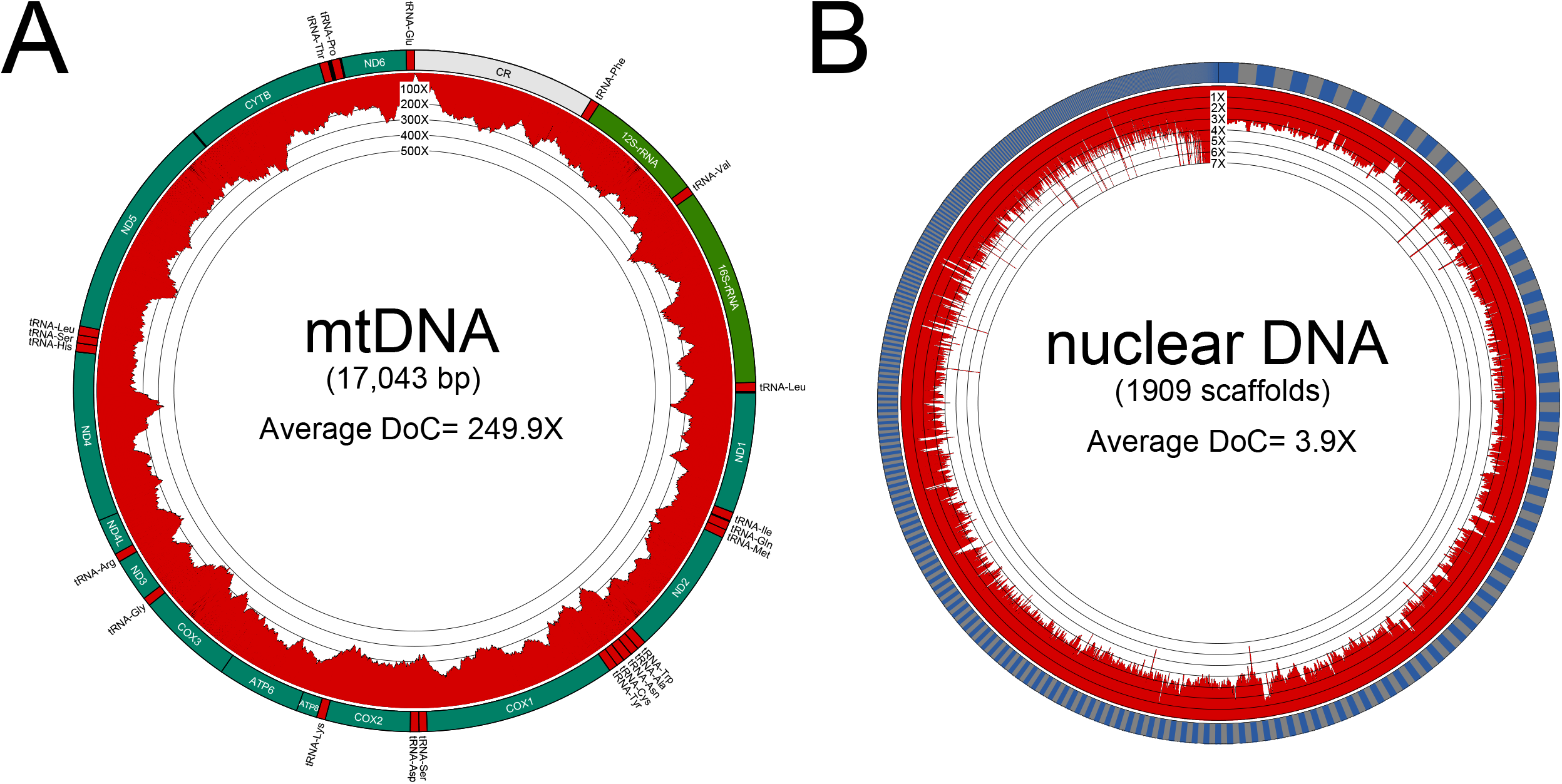
Little bush moa genome assemblies. A) *De novo* assembled mitochondrial genome, with locations of annotated genes and RNAs indicated. Inward-facing plot shows per-base depth of coverage (DoC). B) Reference-based nuclear genome assembly (illustrated for the original moa assembly). Alternating gray and blue sections along outer circle indicate individual scaffolds in order of decreasing size. Inward-facing plot shows depth of coverage calculated in ten nonoverlapping windows per scaffold.

Iterative mapping to a high quality emu reference (Sackton et al. 2019) recovered almost 900 Mb of the little bush moa nuclear genome (GenBank accessions pending), or approximately 75% of the 1.2 Gb emu reference (Fig. 2B, Table 1). Average depth of coverage across called bases was 3.9X for the original assembly, with 87% of bases having DoC ≥ 2 (Fig. 2B, corresponding values for the mapDamage corrected assembly are 4.0X average DoC, with 87% of called bases having DoC ≥ 2). Although moa contigs are relatively short (Table 1, maximum contig length= 12.2 kb for the original assembly and 12.0 kb for mapDamage corrected), the average break between contigs is also small (Table 1, mean contig break= 218 bp for the original assembly and 226 bp for mapDamage corrected). Therefore, more than 85% of BUSCO singlecopy orthologs for birds were identified in moa, with more than 72% of BUSCOs represented by complete coding sequence (Table 1).

### Effect of damage correction on paleognath phylogeny

Sackton et al. (2019) and Cloutier et al. (2019) both used the original (uncorrected) moa genome for phylogenetic analysis. Across all scaffolds, 98% of bases were identical between the original and mapDamage corrected moa genome assemblies, with similarly high sequence identity observed for markers used for phylogenetic analysis (99% of moa bases were identical for conserved non-exonic elements [CNEEs], and 98% of bases were identical for both introns and ultraconserved elements [UCEs]). The recovered sister group to moa was unaffected in most maximum-likelihood gene trees inferred with RAxML (Stamatakis 2014), with 11,698/12,643 (92.5%) of CNEEs, 4,949/4,982 (99.3%) of introns, and 3,133/3,146 (99.6%) of UCEs that had moa sequence recovering the same sister group to moa regardless of whether the original or corrected sequence was used. Futhermore, for the 991 gene trees in total that recovered a different sister group to moa, 522 loci (52.7%) actually had no moa sequence differences and 862 loci (87%) recovered the sister group to moa with bootstrap support below 50%, indicating that observed differences are largely attributable to the random resolution of uninformative gene tree polytomies rather than to well-supported differences caused by mapDamage correction of the moa genome. Species tree inference with the coalescent method MP-EST (Liu et al. 2010) accordingly recovered identical topologies for each marker type and for the total evidence tree combining all loci to that obtained from gene trees using the uncorrected moa sequence, with maximal bootstrap support throughout. We therefore conclude that damage correction of the moa genome has little detectable effect on the phylogeny of paleognathous birds.

### Identification of polymorphic microsatellite markers

Microsatellites offer an appealing option for aDNA studies since these nuclear markers are often highly polymorphic, are spread throughout the genome, and are sufficiently short to allow amplification in degraded samples (Selkoe and Toonen 2006). However, wet-lab approaches for microsatellite isolation are not amenable to degraded aDNA samples, and crossspecies amplification of markers from extant taxa are often unsuccessful (Selkoe and Toonen 2006; Allentoft et al. 2009, 2011). HTS can circumvent these difficulties by identifying microsatellites directly from sequencing reads in the target species. This approach was employed in moa where Allentoft et al. (2009, 2011) developed six polymorphic microsatellites from GS FLX 454 pyrosequencing reads and demonstrated their utility for studies of moa kinship (Allentoft et al. 2015) and population demography (Allentoft et al. 2014).

We used a complementary approach to isolate polymorphic microsatellites from the little bush moa nuclear assembly. We identified 27,127 dinucleotide and 25,170 trinucleotide repeats, approximately half of which met our criteria for inclusion based on flanking sequence contiguity (retaining 14,902 dinucleotides and 13,951 trinucleotides). From these, we identified 40 microsatellites (28 dinucleotides and 12 trinucleotides, Supplementary Table S2) that are heterozygous in the sequenced individual and hence at least minimally polymorphic in the species as a whole. We offer the realigned BAM files for each locus and alignments to other ratites as a community resource for future studies (available in Dryad, an example of each data type is given in Supplementary Fig. S3).

### Tests of selection in candidate genes for limb development

Arguably the most remarkable moa trait is the complete absence of forelimb skeletal elements comprising the wings. All ratites exhibit some degree of forelimb reduction; however, moa are unique in retaining only a fused scapulocoracoid within the pectoral girdle (Worthy and Holdaway 2002; Huynen et al. 2014). Huynen et al. (2014) recovered moa coding sequence for the T-box transcription factor *TBX5,* which plays a key role in forelimb specification and outgrowth (Tanaka 2013; Tickle 2015; Petit et al. 2017), and demonstrated that moa *TBX5* sequence activates promoters of downstream genes in developing chicken embryos. Therefore, alterations to this coding region alone appear unlikely to underlie the wingless moa phenotype (Huynen et al. 2014). We build upon this work by reporting moa sequence for a more comprehensive suite of candidate genes with established involvement in vertebrate limb development (Table 2a), as well as candidates with putative function-altering variants in the Galapagos cormorant hypothesized to accompany forelimb reduction in this flightless species (Burga et al. 2017, Table 2b).

We recovered moa sequence for all investigated genes, with an average 88% of coding sequence per gene recovered from the original moa assembly (Table 2; 87% in the mapDamage corrected version, Supplementary Table S3). We found no frameshift mutations and only a single in-frame stop codon in *HOXD4* which, however, occurred at 1X coverage in both assembly versions and could represent a sequence artifact (this codon was intentionally masked by Ns for further tests). There was no evidence for lineage-specific diversifying selection in moa, with P > 0.05 in aBSREL tests for each gene. RELAX tests also found no significant difference in the strength of selection in moa relative to other ratites for candidate genes with established roles in limb development (Table 2a, Supplementary Table S3a). However, RELAX did identify a significant intensification of selection in the *FAT1* gene in moa and relaxation in *GLI2* relative to other ratites among the 11 candidates originating from study of the Galapagos cormorant (Table 2b, Supplementary Table S3b). Neither of these results remains significant under a more conservative genome-wide correction for an estimated 16,255 genes in birds as opposed to correcting only for the set of 37 candidates tested here (both P > 0.05).

RELAX tests also identified seven candidates with significant selective shifts in flightless lineages relative to other birds (Table 2; 10 candidates, including the previous seven, when mapDamage corrected sequence is used, Supplementary Table S3). Of these, we find evidence for intensified selection in *HOXA2, HOXA4,* and *HOXD4* (additionally *HOXA10, SHH,* and *WNT2B* using the corrected moa sequence), and relaxation in four genes *(GLI3, EVC, FAT1,* and *TALPID3;* note that *FAT1* shows intensified selection in moa relative to other ratites, but also relaxed selection in flightless birds generally). However, only the intensification for *HOXA2* remains significant under the more stringent genome-wide false discovery rate correction (P= 0.021 for both data sets).

PROVEAN analysis identified 24 moa sequence variants of possible functional relevance compared to the emu reference (Supplementary Table S4). Identified sites were identical for both moa assemblies, except *DVL1* where only the alternate moa allele with codon AAC was recovered after base quality recalibration. However, half of these variants (12 of 24) are either shared with other species or are polymorphic in moa, with the emu residue present as an alternative moa allele, indicating that this subset of sites is unlikely to underlie the wingless moa phenotype. Additionally, 16 of the 24 sites display alternative residues in other birds that are often accompanied by PROVEAN scores comparable to moa (Supplementary Table S4). Comparison to an inferred reconstruction of the common ancestor of moa and tinamous yielded broadly similar results, with 17 of 19 potentially functionally relevant moa variants identical to those identified from comparison to the emu reference (Supplementary Table S5).

Putative function-altering variants in the Galapagos cormorant are not shared with other flightless lineages (Supplementary Table S6), indicating that any commonality in the genetic basis for independent losses of flight involving these genes is likely not attributable to convergent or parallel amino acid changes. Burga et al. (2017) also identified a deletion in *CUX1* of the Galapagos cormorant, with experimental assays indicating this gene acts as a transcriptional activator of targets *FAT1* and *OFD1.* As with the other reported Galapagos cormorant variants, the *CUX1* deletion is not shared by moa or other ratites (Supplementary Fig. S4, identical sequence occurs in both moa assemblies). Altogether, we conclude that loss of wings in moa is not attributable to gene loss or pseudogenization within this candidate gene set, although the functional relevance of variants unique to moa requires further experimental work.

## Conclusion

This first nuclear genome assembly begins a new chapter in the already extensive history of moa aDNA research. This genomic resource has already proven useful to assemble genome-wide data sets of nuclear markers for phylogenetic inference (Baker et al. 2014; Cloutier et al. 2019; Sackton et al. 2019). Here, we further demonstrate its utility to isolate markers for population-level studies and to investigate sequence evolution in candidate protein coding genes. The relative contributions of coding sequence variation and mutations in noncoding regulatory elements to phenotypic variation constitute an area of active research (Petit et al. 2017; Lamichhaney et al. 2019; Sackton et al. 2019), and we anticipate that availability of a moa nuclear genome will also contribute to experimental studies of regulatory changes associated with flightless phenotypes.

## Acknowledgements

We thank Oliver Haddrath for extracting moa aDNA, and we thank Trevor Worthy and Paul Scofield for helpful discussion concerning the provenance of the sequenced moa specimen. We thank Sergio Pereira of The Centre for Applied Genomics, The Hospital for Sick Children, Toronto, Canada for overseeing library construction and sequencing. Computations were performed on the GPC supercomputer at the SciNet HPC Consortium funded by Compute Canada, the Government of Ontario, and the University of Toronto, as well as the Odyssey cluster supported by the FAS Division of Science, Research Computing Group at Harvard University. This work was supported by the Natural Science and Engineering Research Council of Canada [to A.J.B.]; the Royal Ontario Museum Governors Fund [to A.J.B.]; and the National Science Foundation [NSF grant DEB 1355343 (EAR 1355292) to A.J.B. and S.V.E.]. This work is dedicated to the memory of coauthor Allan Baker, who was the driving force behind this project but passed away prior to its completion.

---

**Suppl. Fig. S1** Patterns of DNA damage estimated for reads mapping to the little bush moa nuclear (A, B) and mitochondrial (C, D) genomes are consistent with expectations for aDNA. A,C) Excess of purines immediately preceding strand breaks (note that observed coordinates for purine enrichment are shifted slightly relative to expectations likely due to some fragments resulting from DNA shearing during library preparation). B,D) Increased C-to-T transitions towards fragment edge.

**Suppl. Fig. S2** Mitochondrial control region sequence corroborates taxonomic identity of the little bush moa specimen. Sequence from the current study is shown at the top, with representative little bush moa sequences aligned below. GenBank accessions for published sequences are followed by specimen identifiers from Bunce et al. (2009), and dots indicate identity with the first alignment sequence. Samples originating from the South Island of New Zealand are shaded in blue, those from the North Island in yellow, and the 30 bp control region ‘snippet’ of McCallum et al. (2013) is boxed.

**Suppl. Fig. S3** Polymorphic microsatellite repeat identified from the little bush moa nuclear genome. A) Partial sequence for microsatellite anoDid_tri6 showing the consensus genome sequence in black, with mapped reads below. Partners belonging to the same read pair are shaded with the same color, while unpaired reads or those whose partner falls outside the illustrated region are shaded in gray. The microsatellite repeat region is boxed, showing the heterozygous GTT_7_/GTT_9_ genotype for this individual. B) Multiple sequence alignment with the corresponding genomic region from other ratites. The microsatellite repeat region is boxed.

**Suppl. Fig. S4** The 4 amino acid deletion in *CUX1* of the Galapagos cormorant *(P. harrisi)* described by Burga et al. (2017) is not shared with other flightless birds (penguins and ratites).

