## Supplementary figures and images for "First nuclear genome assembly of an extinct moa species, the little bush moa (*Anomalopteryx didiformis*)"

### Supplementary Figures

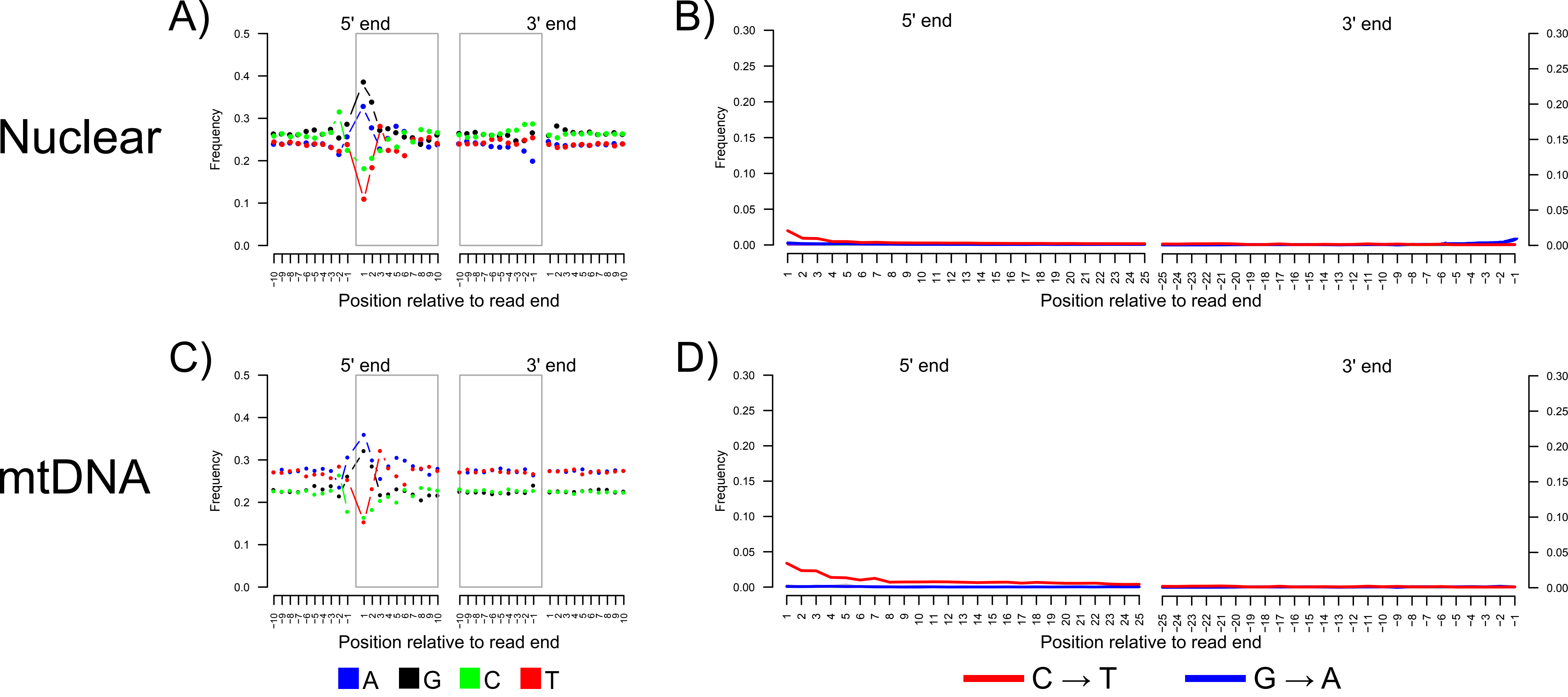

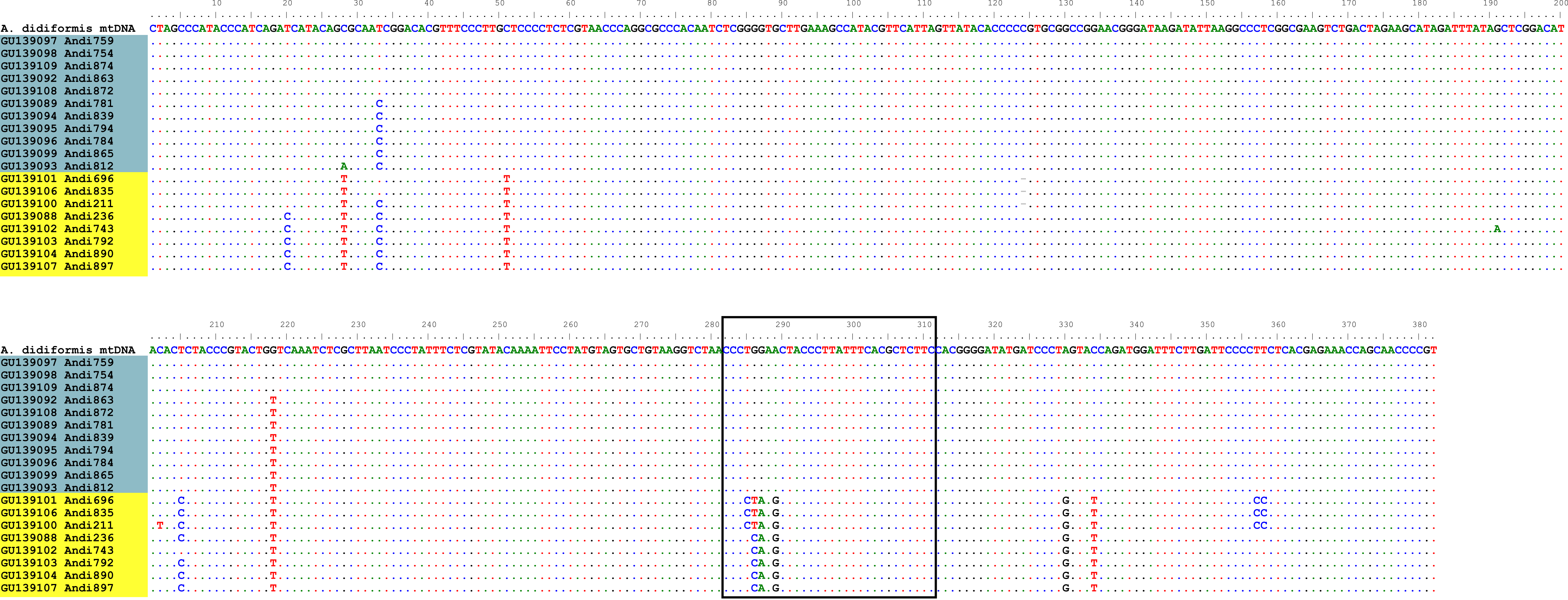

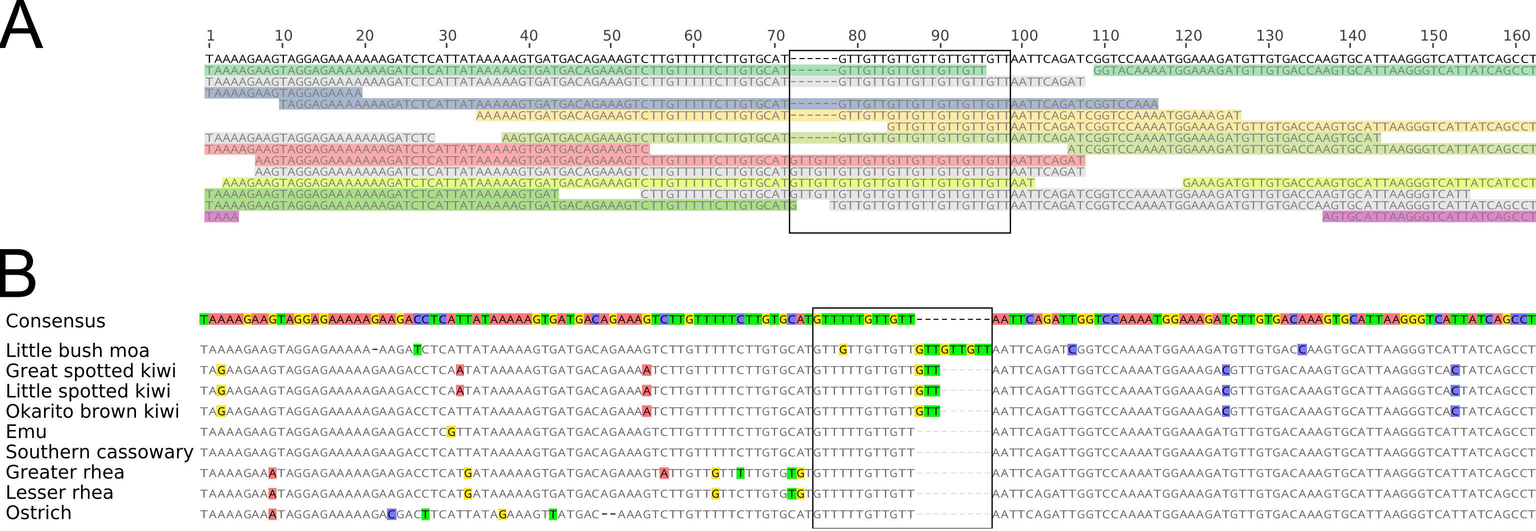

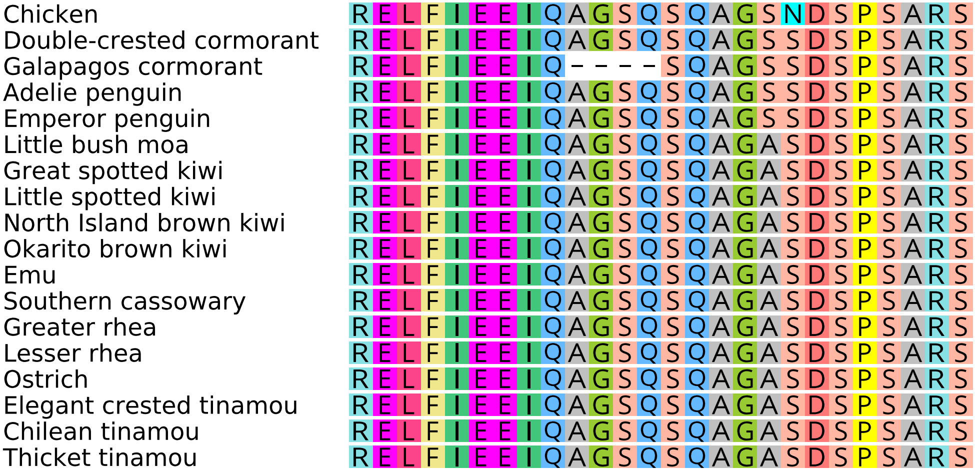
