## Supplementary Tables for "First nuclear genome assembly of an extinct moa species, the little bush moa (*Anomalopteryx didiformis*)"

**Suppl. Table S1** Read mapping information for little bush moa sequencing libraries

| | TruSeq libraries | | Nextera libraries | | |
|---|---|---|---|---|---|
| | **CTTGTA** | **GCCAAT** | **AGGCAG** | **CAGAGA** | **CTCTCT** |
| Raw reads (paired) | 530,099,258 | 59,208,990 | 46,836,756 | 35,552,367 | 38,311,748 |
| Trimmed reads | | | | | |
| Paired | 486,360,442 | 57,743,619 | 45,142,645 | 34,079,453 | 36,434,453 |
| Single | 27,247,969 | 1,059,500 | 44,515 | 35,261 | 46,412 |
| Alignment rate (MAPQ ≥ 30, %) | 12.9 | 9.4 | 0.3 | 0.4 | 0.3 |
| Duplicates (% of mapped) | 69.4 | 98.7 | 75.0 | 81.9 | 73.7 |
| Mean mapped read length (bp)[1] | | | | | |
| Paired | 98.7 | 94.2 | 72.7 | 69.5 | 67.8 |
| Single | 84.2 | 69.7 | 86.7 | 63.0 | 49.6 |
| Mean insert size (bp)[1] | 206.8 | 274.0 | 90.4 | 78.6 | 74.9 |

[1]for de-duplicated reads

**Suppl. Table S2** Polymorphic di- and trinucleotide microsatellite repeats identified in little bush moa

| Locus | Scaffold | Start (Repeat)[1] | End (Repeat)[1] | Start (Repeat + flank)[1] | End (Repeat + flank)[1] | REF Allele | ALT Allele |
|---|---|---|---|---|---|---|---|
| **Dinucleotide repeats** | | | | | | | |
| anoDid_di1 | scaffold_423 | 460927 | 460943 | 460680 | 461121 | $(AG)_8$ | $(AG)_9$ |
| anoDid_di2 | scaffold_48 | 3097112 | 3097126 | 3096862 | 3097375 | $(GT)_7$ | $(GT)_6$ |
| anoDid_di3 | scaffold_50 | 996879 | 996901 | 996641 | 997129 | $(AG)_{11}$ | $(AG)_9$ |
| anoDid_di4 | scaffold_6 | 2351922 | 2351934 | 2351691 | 2352158 | $(CT)_6$ | $(CT)_5$ |
| anoDid_di5 | scaffold_60 | 2709332 | 2709358 | 2709113 | 2709584 | $(GT)_{13}$ | $(GT)_{12}$ |
| anoDid_di6 | scaffold_79 | 2063607 | 2063619 | 2063357 | 2063853 | $(CT)_6$ | $(CT)_7$ |
| anoDid_di7 | scaffold_81 | 3721969 | 3721991 | 3721739 | 3722214 | $(GT)_{11}$ | $(GT)_{13}$ |
| anoDid_di8 | scaffold_126 | 739242 | 739262 | 739086 | 739499 | $(GT)_{10}$ | $(GT)_{11}$ |
| anoDid_di9 | scaffold_135 | 108054 | 108070 | 107877 | 108297 | $(CT)_8$ | $(CT)_9$ |
| anoDid_di10 | scaffold_136 | 1740662 | 1740678 | 1740422 | 1740901 | $(AC)_8$ | $(AC)_7$ |
| anoDid_di11 | scaffold_158 | 2414896 | 2414908 | 2414745 | 2415138 | $(AC)_6$ | $(AC)_7$ |
| anoDid_di12 | scaffold_17 | 5603622 | 5603638 | 5603392 | 5603884 | $(AC)_8$ | $(AC)_9$ |
| anoDid_di13 | scaffold_18 | 221485 | 221501 | 221325 | 221741 | $(AG)_8$ | $(AG)_7$ |
| anoDid_di14 | scaffold_202 | 427690 | 427702 | 427459 | 427936 | $(AC)_6$ | $(AC)_5$ |
| anoDid_di15 | scaffold_21 | 4341655 | 4341667 | 4341405 | 4341898 | $(CT)_6$ | $(CT)_7$ |
| anoDid_di16 | scaffold_225 | 1492187 | 1492211 | 1491957 | 1492454 | $(AC)_{12}$ | $(AC)_{11}$ |
| anoDid_di17 | scaffold_26 | 3615436 | 3615450 | 3615225 | 3615686 | $(AG)_7$ | $(AG)_6$ |
| anoDid_di18 | scaffold_26 | 5268739 | 5268753 | 5268489 | 5268983 | $(AC)_7$ | $(AC)_9$ |
| anoDid_di19 | scaffold_105 | 1805044 | 1805062 | 1804941 | 1805300 | $(AG)_9$ | $(AG)_8$ |
| anoDid_di20 | scaffold_28 | 3056771 | 3056783 | 3056537 | 3056994 | $(AG)_6$ | $(AG)_7$ |
| anoDid_di21 | scaffold_30 | 1241412 | 1241432 | 1241184 | 1241551 | $(GT)_{10}$ | $(GT)_{11}$ |
| anoDid_di22 | scaffold_30 | 3716948 | 3716962 | 3716704 | 3717204 | $(AT)_7$ | $(AT)_6$ |
| anoDid_di23 | scaffold_107 | 1738937 | 1738949 | 1738688 | 1739195 | $(CT)_6$ | $(CT)_7$ |
| anoDid_di24 | scaffold_317 | 517054 | 517070 | 516817 | 517315 | $(AC)_8$ | $(AC)_6$ |
| anoDid_di25 | scaffold_368 | 665439 | 665465 | 665244 | 665711 | $(GT)_{13}$ | $(GT)_{14}$ |
| anoDid_di26 | scaffold_37 | 4246825 | 4246843 | 4246575 | 4246957 | $(AG)_9$ | $(AG)_8$ |
| anoDid_di27 | scaffold_40 | 4006873 | 4006887 | 4006627 | 4007126 | $(AT)_7$ | $(AT)_6$ |
| anoDid_di28 | scaffold_41 | 1770175 | 1770189 | 1769925 | 1770423 | $(GT)_7$ | $(GT)_6$ |
| **Trinucleotide repeats** | | | | | | | |
| anoDid_tri1 | scaffold_494 | 312051 | 312078 | 311833 | 312325 | $(AAC)_9$ | $(AAC)_8$ |
| anoDid_tri2 | scaffold_67 | 2212411 | 2212423 | 2212166 | 2212623 | $(GCT)_4$ | $(GCT)_5$ |
| anoDid_tri3 | scaffold_84 | 2470005 | 2470020 | 2469771 | 2470233 | $(AGC)_5$ | $(AGC)_6$ |
| anoDid_tri4 | scaffold_91 | 3565159 | 3565171 | 3564951 | 3565405 | $(GTT)_4$ | $(GTT)_5$ |
| anoDid_tri5 | scaffold_124 | 1297756 | 1297768 | 1297686 | 1298004 | $(AGG)_4$ | $(AGG)_3$ |
| anoDid_tri6 | scaffold_151 | 866522 | 866543 | 866278 | 866655 | $(GTT)_7$ | $(GTT)_9$ |
| anoDid_tri7 | scaffold_219 | 836843 | 836867 | 836607 | 837000 | $(CCT)_8$ | $(CCT)_9$ |
| anoDid_tri8 | scaffold_239 | 709586 | 709616 | 709338 | 709856 | $(CGG)_{11}^{\dagger}$ | $(CGG)_8$ |
| anoDid_tri9 | scaffold_309 | 618407 | 618419 | 618184 | 618614 | $(AAG)_4$ | $(AAG)_5$ |
| anoDid_tri10 | scaffold_320 | 505222 | 505234 | 505000 | 505455 | $(GCT)_4$ | $(GCT)_5$ |
| anoDid_tri11 | scaffold_351 | 134678 | 134693 | 134458 | 134897 | $(GCT)_5$ | $(GCT)_4$ |
| anoDid_tri12 | scaffold_36 | 1888459 | 1888474 | 1888223 | 1888661 | $(GCT)_6^{\dagger}$ | $(GCT)_7$ |

† differs from genome assembly by +1 repeat unit following indel realignment

[1] Coordinates are given for the original moa assembly. Refer to the accompanying Dryad Digital Repository archive for the corresponding positions in the mapDamage corrected assembly.

**Suppl. Table S3** Tests of selection for candidate limb development genes using moa sequence from the mapDamage corrected genome assembly

| Gene | Description | CDS length (AA, % of total) | | | RELAX tests | | | |
|---|---|---|---|---|---|---|---|---|
| | | | | | Moa | | All flightless | |
| | | Chicken | Emu | Moa | K | $P_{adj}$ | K | $P_{adj}$ |
| **a) Candidate limb development genes** | | | | | | | | |
| FGF8 | Fibroblast growth factor 8 | 214 | 214 | 188 (88%) | 3.931 | 0.991 | 49.895 | 0.057 |
| FGF10 | Fibroblast growth factor 10 | 212 | 212 | 180 (85%) | 0.495 | 0.464 | 1.052 | 0.438 |
| GLI3 | GLI family zinc finger 3 | 1576 | 1575 | 1568 (99%) | 1.464 | 0.416 | 0.358 | 0.004 |
| HOXA1 | Homeobox A1 | 320 | 319 | 288 (90%) | 1.046 | 0.973 | 0.947 | 0.410 |
| HOXA2 | Homeobox A2 | 375 | 374 | 358 (96%) | 0.177 | 0.074 | 3.414 | < 0.001 |
| HOXA3 | Homeobox A3 | 413 | 413 | 414 (100%) | 1.491 | 0.416 | 1.033 | 0.424 |
| HOXA4 | Homeobox A4 | 309 | 145† | 155 (50%) | 0.365 | 0.416 | 2.564 | 0.004 |
| HOXA5 | Homeobox A5 | 270 | 270 | 251 (93%) | 29.348 | 0.289 | 0.633 | 0.172 |
| HOXA6 | Homeobox A6 | 231 | 231 | 231 (100%) | 0.828 | 0.808 | 0.272 | 0.292 |
| HOXA7 | Homeobox A7 | 219 | 219 | 216 (99%) | 32.541 | 0.416 | 1.157 | 0.340 |
| HOXA9 | Homeobox A9 | 260 | 261 | 249 (95%) | 1.985 | 0.416 | 2.040 | 0.060 |
| HOXA10 | Homeobox A10 | 364 | 317† | 289 (79%) | 1.239 | 0.934 | 2.914 | 0.041 |
| HOXA11 | Homeobox A11 | 297 | 297 | 254 (86%) | 0.305 | 0.416 | 1.465 | 0.079 |
| HOXA13 | Homeobox A13 | 290 | 290 | 269 (93%) | 1.179 | 0.934 | 0.853 | 0.382 |
| HOXD3 | Homeobox D3 | 413 | 248† | 247 (60%) | 1.211 | 0.808 | 1.326 | 0.340 |
| HOXD4 | Homeobox D4 | 237 | 237 | 200 (85%) | 1.123 | 0.970 | 9.628 | 0.004 |
| HOXD8 | Homeobox D8 | 268 | 147† | 146 (54%) | 0.607 | 0.517 | < 0.001 | 0.304 |
| HOXD9 | Homeobox D9 | 302 | 299 | 283 (94%) | 0.295 | 0.365 | 0.816 | 0.173 |
| HOXD10 | Homeobox D10 | 339 | 339 | 339 (100%) | 0.923 | 0.991 | 49.998 | 0.121 |
| HOXD11 | Homeobox D11 | 280 | 282 | 272 (96%) | 0.967 | 0.973 | 0.642 | 0.057 |
| HOXD12 | Homeobox D12 | 266 | 266 | 266 (100%) | 1.452 | 0.416 | 1.006 | 0.474 |
| HOXD13 | Homeobox D13 | 301 | 82† | 74 (25%) | 0.926 | 0.991 | 0.939 | 0.424 |
| SALL4 | Spalt-like transcription factor 4 | 1108 | 1111 | 1023 (92%) | 0.793 | 0.416 | 0.858 | 0.113 |
| SHH | Sonic hedgehog | 425 | 422 | 396 (94%) | 0.772 | 0.808 | 2.011 | 0.001 |
| TBX5 | T-box 5 | 521 | 538 | 419 (78%) | 0.243 | 0.416 | 1.024 | 0.450 |
| WNT2B | Wnt family member 2B | 385 | 330† | 257 (67%) | 0.769 | 0.517 | 1.748 | 0.014 |
| **b) Candidate genes from the Galapagos cormorant** | | | | | | | | |
| DCHS1 | Dachsous cadherin-related 1 | 3266 | 3267 | 3072 (94%) | 1.098 | 0.416 | 1.007 | 0.451 |
| DVL1 | Dishevelled segment polarity protein 1 | 712 | 655† | 633 (89%) | 1.430 | 0.416 | 0.419 | 0.079 |
| DYNC2H1 | Dynein cytoplasmic 2 heavy chain 1 | 4301 | 4295 | 3968 (92%) | 1.137 | 0.416 | 0.755 | 0.060 |
| EVC | EvC ciliary complex subunit 1 | 984 | 927† | 870 (88%) | 0.168 | 0.416 | 0.642 | 0.014 |
| FAT1 | FAT atypical cadherin 1 | 4645 | 4644 | 4473 (96%) | 1.592 | 0.013 | 0.878 | 0.014 |
| GLI2 | GLI family zinc finger 2 | 1528 | 1528 | 1527 (100%) | 0.405 | 0.013 | 0.928 | 0.212 |
| IFT122 | Intraflagellar transport 122 | 1245 | 1239 | 1180 (95%) | 1.126 | 0.621 | 1.355 | 0.113 |
| KIF7 | Kinesin family member 7 | 1412 | 1279† | 1225 (87%) | 0.845 | 0.416 | 35.383 | 0.057 |
| OFD1 | OFD1, centriole and centriolar satellite protein | 1012 | 1014 | 971 (96%) | 0.986 | 0.991 | 0.153 | 0.079 |
| TALPID3 | KIAA0586 | 1523 | 1527 | 1432 (94%) | 0.766 | 0.416 | 0.035 | 0.014 |
| WDR34 | WD repeat domain 34 | 500 | 502 | 459 (91%) | 18.909 | 0.251 | 0.776 | 0.077 |

K: Relaxation parameter (values < 1 indicate relaxed selection on foreground branches, values > 1 denote intensified selection)
$P_{adj}$: Adjusted P-value (Q-value) controlling for the false discovery rate at a significance level of 0.05 based on N= 37 genes tested
†Partial CDS recovered in the emu reference sequence

**Suppl. Table S4** Moa variants with PROVEAN score < -5 compared to the emu reference

| Gene | Variant[1] | Alignment Position (AA)[2] | PROVEAN (Emu-Moa) | Shared[3] | Alternative (PROVEAN)[4] | DoC[5] | Moa alleles |
|---|---|---|---|---|---|---|---|
| DCHS1 | P2594L | 2654 | -7.364 | No | S (-5.572) | 3X | |
| | Q3065del | 3126 | -5.600 | Yes | P (-1.429) | 6X | |
| | H3199L | 3267 | -6.496 | Yes | n/a | 1X | |
| DVL1 | N206Y | 263 | -5.080 | No | n/a | 2X | Tyr Y (TAC, 1X DoC)<br>Asn N (AAC, 1X DoC) |
| DYNC2H1 | P789A | 793 | -6.517 | No | n/a | 2X | |
| | M951T | 955 | -5.099 | No | n/a | 2X | |
| | C2494Y | 2498 | -6.584 | No | R (-5.204) | 6X | |
| EVC | H544R | 603 | -5.103 | Yes | Q (-4.455) | 4X | |
| | D703G | 762 | -5.620 | Yes | E (-3.157)<br>N (-4.149) | 6X | |
| FAT1 | E2568V | 2570 | -5.223 | No | G (-4.742)<br>K (-2.900)<br>Q (-2.032) | 1X | |
| | K2919I | 2921 | -6.160 | No | R (-2.203) | 4X | |
| | P3411R | 3413 | -6.287 | Yes | A (-4.538)<br>S (-5.030) | 5X | |
| | H4379P | 4385 | -5.295 | No | N (-3.224)<br>Y (-2.563) | 2X | |
| GLI2 | A951_L952insL | 982 | -7.986 | No | n/a | 6X | |
| HOXD8 | E124G | 258 | -6.488 | No | n/a | 4X | |
| KIF7 | L459Q | 641 | -5.210 | Yes | M (-1.685) | 6X | |
| OFD1 | E190A | 190 | -5.926 | No | K (-3.947)<br>Q (-2.958) | 6X | |
| | R896del | 917 | -6.903 | Yes | K (-1.404)<br>M (-3.376)<br>S (-3.318) | 5X | |
| TALPID3 | H766Q | 856 | -7.407 | Yes | n/a | 3X | Gln Q (CAG, 2X DoC)<br>His H (CAC, 1X DoC) |
| | P817A | 907 | -7.738 | Yes | T (-7.691) | 5X | |
| | P1163A | 1281 | -7.066 | Yes | n/a | 7X | |
| | P1218A | 1336 | -6.453 | Yes | H (-6.836)<br>L (-7.451)<br>S (-6.270)<br>T (-6.366) | 6X | |
| | P1229R | 1347 | -6.538 | No | A (-6.168) | 5X | |
| | P1379L | 1501 | -6.853 | No | A (-5.107)<br>S (-5.243) | 7X | |

[1]Variants are listed using HGVS (Human Genome Variation Society) notation. For example, P2594L indicates P at position 2594 in the emu reference amino acid sequence is replaced by L in moa.
[2]Numbering refers to column in the amino acid alignment of all species provided in the accompanying Dryad data release.
[3]Indicates whether moa amino acid replacement is shared by other birds in alignment.
[4]Indicates alternative amino acid replacement present in other birds in alignment, with PROVEAN score relative to the emu reference in brackets.
[5]Depth of coverage (DoC) is given for the original moa assembly, using de-duplicated reads and with overlapping read pairs counted as 1X coverage.

**Suppl. Table S5** Moa variants with PROVEAN score < -5 compared to a moa-tinamou ancestral reference sequence. Variants differing from those using an emu reference are shown in bold.

| Gene | Variant[1] | Alignment Position (AA)[2] | PROVEAN (Anc-Moa) | Emu reference equivalent | Shared[3] | Alternative (PROVEAN)[4] | DoC[5] | Moa alleles |
|---|---|---|---|---|---|---|---|---|
| DCHS1 | **P1989L** | **1989** | **-5.116** | **P1945L (-4.799)** | **No** | **S (-2.920)** | **5X** | |
| | P2654L | 2654 | -7.344 | P2594L (-7.364) | No | S (-5.572) | 3X | |
| | Q3126del | 3126 | -5.064 | Q3065del (-5.600) | Yes | P (-1.429) | 6X | |
| DVL1 | N263Y | 263 | -5.069 | N206Y (-5.080) | No | n/a | 2X | Tyr Y (TAC, 1X DoC)<br>Asn N (AAC, 1X DoC) |
| DYNC2H1 | P793A | 793 | -6.518 | P789A (-6.514) | No | n/a | 2X | |
| | M955T | 955 | -5.233 | M951T (-5.099) | No | n/a | 2X | |
| | C2498Y | 2498 | -6.980 | C2494Y (-6.584) | No | R (-5.204) | 6X | |
| EVC | H603R | 603 | -5.087 | H544R (-5.103) | Yes | Q (-4.455) | 4X | |
| | D762G | 762 | -5.516 | D703G (-5.620) | Yes | E (-3.157)<br>N (-4.149) | 6X | |
| FAT1 | E2570V | 2570 | -5.409 | E2568V (-5.223) | No | G (-4.742)<br>K (-2.900)<br>Q (-2.032) | 1X | |
| | K2921I | 2921 | -6.226 | K2919I (-6.160) | No | R (-2.203) | 4X | |
| | H4385P | 4385 | -5.216 | H4379P (-5.295) | No | N (-3.224)<br>Y (-2.563) | 2X | |
| HOXD8 | E258G | 258 | -6.549 | E124G (-6.488) | No | n/a | 4X | |
| OFD1 | E190A | 190 | -5.961 | E190A (-5.926) | No | K (-3.947)<br>Q (-2.958) | 6X | |
| TALPID3 | H856Q | 856 | -7.335 | H766Q (-7.407) | Yes | n/a | 3X | Gln Q (CAG, 2X DoC)<br>His H (CAC, 1X DoC) |
| | **M1127T** | **1127** | **-5.056** | **M1028T (-4.731)** | **No** | **I (-3.099)<br>L (-2.436)<br>V (-3.059)** | **8X** | **Thr T (ACG, 7X DoC)<br>Thr T (ACA, 1X DoC)** |
| | P1281A | 1281 | -6.827 | P1163A (-7.066) | Yes | n/a | 7X | |
| | P1336A | 1336 | -6.456 | P1218A (-6.453) | Yes | H (-6.836) | 6X | |
| | P1347R | 1347 | -6.440 | P1229R (-6.538) | No | A (-6.168) | 5X | |

[1]Variants are listed using HGVS (Human Genome Variation Society) notation. For example, P1989L indicates P at position 1989 in the moa-tinamou ancestral reference amino acid sequence is replaced by L in moa.
[2]Numbering refers to column in the amino acid alignment of all species provided in the accompanying Dryad data release.
[3]Indicates whether moa amino acid replacement is shared by other birds in alignment.
[4]Indicates alternative amino acid replacement present in other birds in alignment, with PROVEAN score relative to the moa-tinamou ancestor reference in brackets.
[5]Depth of coverage (DoC) is given for the original moa assembly, using de-duplicated reads and with overlapping read pairs counted as 1X coverage.

**Suppl. Table S6** Putative function-altering variants in the Galapagos cormorant (*P. harrisi*) are not shared with other flightless lineages

| Gene | *P. harrisi* residue | Alignment position (AA)[1] | Amino acid | | | | |
|---|---|---|---|---|---|---|---|
| | | | *P. harrisi* | Moa | Other ratites | Penguins | Flighted birds |
| DCHS1 | 2063 | 2107 | D | G | G | G | G |
| DVL1 | 103 | 103 | L | P | P | n/a | P |
| DYNC2H1 | 2733 | 2735 | S | P | P | P | P |
| EVC | 341 | 343 | I | T | T | n/a | T |
| FAT1 | 1717 | 1742 | L | S | S | S | S |
| | 2462 | 2487 | C | Y | Y | Y | Y |
| GLI2 | 1086 | 1127 | T | P | P | P/S | P/S |
| IFT122 | 691 | 924 | L | Q | Q | Q | Q |
| KIF7 | 833 | 965 | W | R | R | R | R |
| OFD1 | 325 | 326 | C | R | R | R | R/C |
| | 517 | 518 | T | K | K | K | K/E |
| | 899 | 924 | G | E | E | E | E/N |
| TALPID3 | 758 | 1005 | V | D | D | D | D/N |
| WDR34 | 188 | 190 | R | P | P | n/a | P |

[1]Numbering refers to column in the amino acid alignment of all species provided in the accompanying Dryad data release
